# Quantitative assessment of cell fate decision between autophagy and apoptosis

**DOI:** 10.1101/129767

**Authors:** Bing Liu, Zoltán N. Oltvai, Hulya Bayir, Gary A. Silverman, Stephen C. Pak, David H. Perlmutter, Ivet Bahar

## Abstract

Autophagy and apoptosis regulate cell survival and death, and are implicated in the pathogenesis of many diseases. The same type of stress signals can induce either process, but it is unclear how cells ‘assess’ cellular damage and make a ‘life’ or ‘death’ decision by activating autophagy or apoptosis. A computational model of coupled apoptosis and autophagy is built here to study the systems-level dynamics of the underlying signaling network. The model explains the differential dynamics of autophagy and apoptosis in response to various experimental stress signals. Autophagic response dominates at low-to-moderate stress; whereas the response shifts from autophagy (graded activation) to apoptosis (switch-like activation) with increasing intensity of stress. The model reveals that this dynamic cell fate decision is conferred by a core regulatory network involving cytoplasmic Ca^2+^ as a rheostat that fine-tunes autophagic and apoptotic responses. A G-protein signaling-mediated feedback loop maintains cytoplasmic Ca^2+^ level, which in turn governs autophagic response through an AMP-activated protein kinase (AMPK)-mediated feedforward loop. The model identified Ca^2+^/calmodulin-dependent kinase kinase β (CaMKKβ) as a determinant of the opposite roles of cytoplasmic Ca^2+^ in autophagy regulation. The results also demonstrated that the model could contribute to the development of pharmacological strategies modulate cell fate decisions.

## Introduction

Autophagy is a cytoprotective homeostatic process in which cells digest their own cytoplasmic constituents and organelles and degrade them in the lysosomes in response to diverse stress stimuli^1^. The resulting products can be recycled to generate energy and build new proteins, hence the activation of autophagy as a protective mechanism against starvation^2^. Autophagy is also a cellular quality control process that removes damaged organelles and aggregates of misfolded proteins that may cause e.g., neurodegenerative diseases^3^ and liver diseases^4,5^. However, excessive autophagy has been linked to cell death, and autophagy activation may be harmful in certain disease conditions (e.g. cancer). The modulation of autophagy has thus emerged as an important therapeutic goal for diverse diseases^6,7^.

Due to a complex crosstalk between autophagy and apoptosis^8^, it is often unclear which specific processes contribute to pro-survival or pro-death effects in a given disease. Previous work suggested that excessive autophagy can induce “autophagic” cell death^8^. Recent studies, however, suggest instead that in dying cells autophagy represents an attempt to prevent the inevitable demise of the cell^9^.

A database resource has developed for systems-level autophagy research^10^; however, it provided static information of the regulatory network implicated in autophagy. Here, we developed a mathematical model to quantitatively assess how cells orchestrate the dynamics of signaling networks to make ‘life’ vs. ‘death’ decisions, and how these are modulated by pharmacological interventions. Our model includes mTOR and inositol signaling autophagic pathways and intrinsic apoptosis pathways as well as their crosstalks mediated by Bcl2, caspases, p53, calpain and Ca^2+^. Using a statistical model checking (SMC)-based framework^11^, we generated a calibrated model that captures cellular heterogeneity and closely reproduces the differential initiation and time evolution of autophagy or apoptosis in response to nutritional, genotoxic, or endoplasmic reticulum (ER) stresses observed in single-cell experiments^12^.

The model points to AMP-activated protein kinase (AMPK) as a key regulator that mediates the competing roles of intracellular Ca^2+^ (Ca^2+^(IC)). The Ca^2+^(IC) level, [Ca^2+^(IC)] acts as a rheostat that fine-tunes autophagic and apoptotic responses, regulated by a positive (G-protein signaling) feedback loop and Ca^2+^/calmodulin-dependent kinase kinase β (CaMKKβ) levels. The model allows for rapid assessment of the effect of a series of drugs on the onset and development of autophagy or apoptosis, under different conditions.

## Results

### Quantitative model of coupled autophagy and apoptosis signaling network

We built a mathematical model that includes the major signaling cascades activated in response to nutritional, genotoxic, and ER stresses (**Fig. 1a**). The model consists of 97 ordinary differential equations (ODEs). Supplementary **Tables S1-S2** list the components (and acronyms), rate equations and parameters. The corresponding reaction network is composed of five modules (**Fig. 1b**):

**Figure 1.**
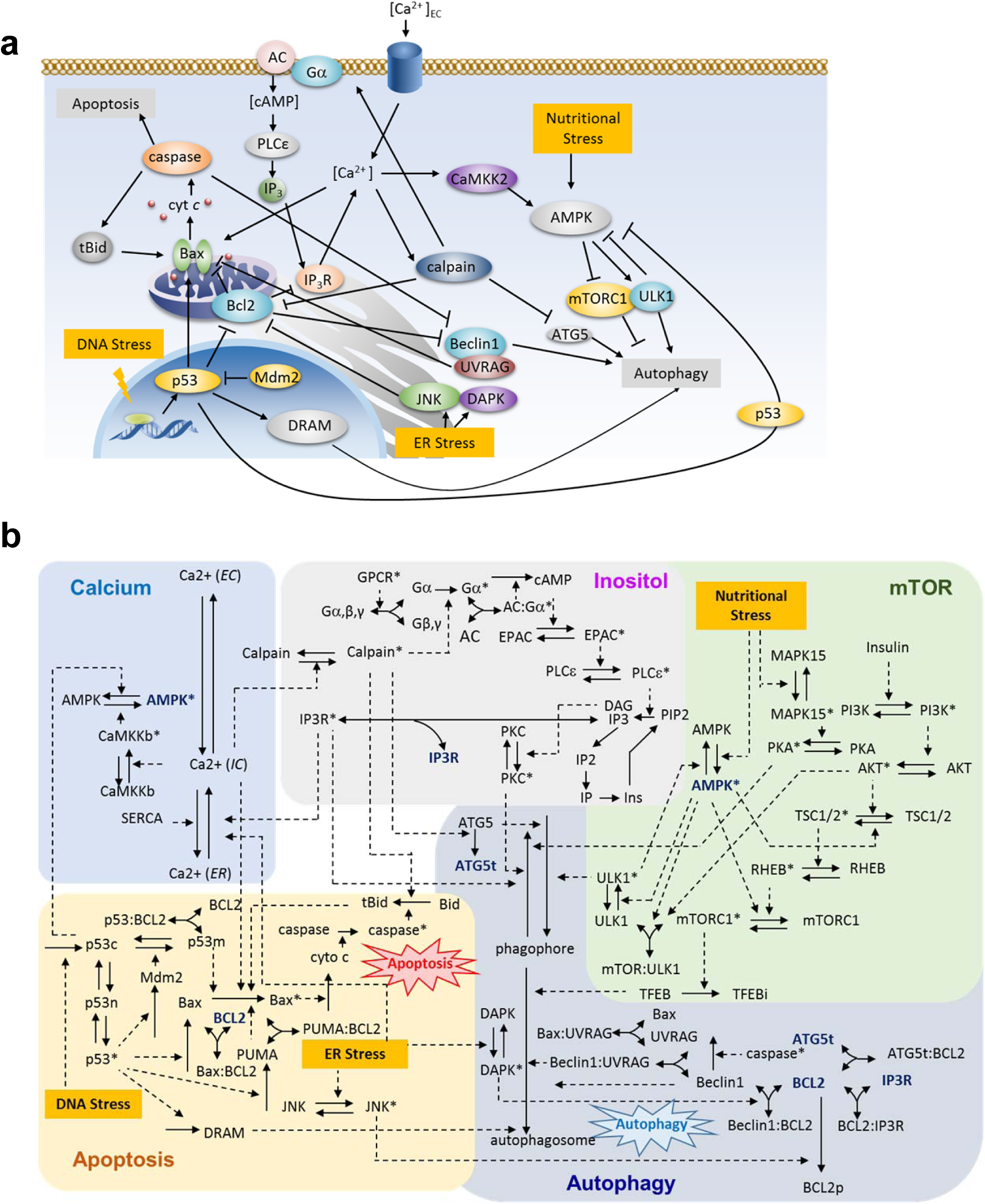
Reaction network model for autophagy-apoptosis crosstalk. (**a**). Schematic illustration of the main components and their key interactions. Activating and inhibitory interactions are distinguished by different types of arrows. Full names of compounds are given in Supplementary **Table S1**. (**b**) A more detailed diagram depicting the network of PPIs. The network is composed of five coupled modules (calcium, inositol, mTOR, apoptosis and autophagy). *Solid* and *dashed* arrows refer to association/disassociation/translocation and chemical reactions, respectively. The complete list of reactions/interactions is given in the Supplementary **Table S2**. Some components involved in multiple modules (e.g. AMPK, IP_3_R, Bcl-2, Bax, Atg5) are shown at multiple places, for clarity.

#### Apoptosis module

Following our previous model^13^, nuclear p53 gains transcriptional activity for pro-apoptotic proteins represented by Bax and its activator, PUMA, under genotoxic (DNA) stress. Activated p53 induces Mdm2, which inhibits p53 by facilitating its ubiquitinylation and translocation to the mitochondria. Mitochondrial p53 further inhibits the anti-apoptotic protein Bcl-2, and activates Bax to promote the formation of mitochondrial outer membrane permeability pores which enables the release of cytochrome *c* (cyt *c*), and leads to caspase activation. The process is amplified by a positive feedback loop involving caspases, Bid, and Bax, in which caspase-truncated Bid (tBid) induces the activation and subsequent oligomerization of Bax. ER stress can also trigger apoptosis, for example, through activation of the JNK- and death-associated protein kinase 1 (DAPK1)-dependent pathways^14^. Further activation of caspase cascades by calpain due to the stress-induced release of ER Ca^2+^ to the cytoplasm is described in the calcium module.

#### Autophagy module

Autophagy involves the formation of phagophore, which engulfs dysfunctional substrates, aggregates, or organelles to form autophagosomes. The content of the autophagosome is degraded through the lysosomal machinery^3^. Autophagic elimination involves several protein complexes such as the mTOR complex 1 (mTORC1) and the Unc-51-like autophagy-activating kinase 1 (ULK1). Our model also includes mediators of autophagy progression such as Atg5, Beclin-1 and UV radiation-resistance associated gene (UVRAG) protein and regulatory proteins (e.g. PKA and PKC) that inhibit autophagy by phosphorylating LC3^15^.

The above two processes are coupled in multiple ways: (i) UVRAG inhibits Bax, while it interacts with Beclin-1 to promote autophagy^16^; (ii) Bcl-2 inhibits autophagy by interacting with Beclin-1^17^, and can be suppressed by truncated Atg5^18^; (iii) Bcl-2 also enhances autophagy via its interaction with inositol 1,4,5 trisphosphate receptor (IP_3_R), an inhibitor of autophagosome formation^19^; (iii) Activated caspases can cleave Beclin-1 to inhibit autophagy^20^; (iv) Cytoplasmic p53 inhibits autophagy by deactivating AMPK^21^, while nuclear p53 promotes autophagy via transcriptional activation of damage-regulated autophagy modulator (DRAM) - a lysosomal protein that induces autophagy^22^, and stimulation of JNK signaling pathways to trigger Bcl-2 phosphorylation^23^; (v) ER stress activates JNK, which, in turn, phosphorylates (and inactivates) Bcl-2^24^; it also activates DAPK, which dissociates from Bcl-2:Beclin-1 complex^25^; (vi) Activated calpain cleaves Atg5 to inhibit autophagy, and Bid to induce apoptosis^18^.

#### mTOR module

Under normal condition, mTORC1 is phosphorylated and active (designated with superscript*); mTORC1* binds ULK1 thus preventing its activation, and inactivates the transcription factor EB (TFEB)^6^, which are essential proteins promoting autophagy. Under cellular stress, stimulation of PI3K-AKT-TSC1/2-RHEB pathway inactivates mTORC1*, leading to the release of ULK1 and activation of autophagy^15^. In parallel, nutrient stress is sensed by AMPK, which inhibits mTORC1 pathways as a mechanism for suppressing cell growth and biosynthesis^26^. Specifically, AMPK releases and thus activates ULK1 which induces autophagy. It also inactivates mTORC1* by triggering the TSC1/2- RHEB cascade and directly phosphorylating a protein (Raptor) in mTORC1^15^. Furthermore, AMPK is negatively regulated by cytoplasmic p53 and ULK1* and positively regulated by CaMKKβ (see the calcium module)^27^.

#### Inositol module

The module is activated upon ligand-binding to G-protein coupled receptors (GPCRs), which prompts the dissociation of the α-subunit of the intracellularly bound G protein from the β- and γ-subunits. Dissociation of activated Gα*s* subtype, Gα*, stimulates the production of cyclic AMP (cAMP) upon binding onto and activating adenylate cyclase (AC) that catalyzes the conversion of ATP to cAMP. The effects of Gα subtypes Gαq and Gαi on AC are implicitly included through model parameters. cAMP blocks autophagy by activating the exchange protein EPAC which in turn activates the phospholipase Cε (PLCε). PLCε* induces the production of IP_3_ and consequently, the release of Ca^2+^ from ER upon binding of IP_3_ to its receptor IP_3_R, a ligand-gated Ca^2+^ channel, on the ER membrane^6^.

#### Calcium module

Ca^2+^ translocates between the extracellular (EC) space, the cytoplasm and the ER, regulated by voltage-gated and ligand-gated ion channels (e.g. IP_3_R) and pumps (e.g. SERCA)^28^. Ca^2+^(IC) activates CaMKKβ, which phosphorylates (or activates) AMPK to promote autophagy^29^ (see mTOR module). CaMKKβ also activates calpain. Calpain* activates the inositol pathway, inhibits autophagy (by cleaving Atg5)^18^, and/or induce apoptosis by activating Bax^30^.

### The calibrated model reproduces differential dynamics of autophagy and apoptosis in response to nutritional-, genotoxic-, and ER-stress

To estimate the unknown parameters, we utilized single-cell-based experimental data^12^ on the time courses of autophagy and apoptosis observed in H4 cells under: (i) 10, 40, 80, and 200 nM Torin 1 (or rapamycin, mTOR inhibitor) treatment, (ii) 0.02, 0.08, 0.32 and 2.5 μM staurosporine (STS, cytotoxic reagent that inhibits several kinases including PKC and PKA) treatment. The resulting kinetic parameters are listed in Supplementary **Table S2**. **Figure 2a-b** display the good agreement achieved between the profiles generated by our simulations with the optimized set of parameters (*dashed curves*) and the experimental data (*dots*).

**Figure 2.**
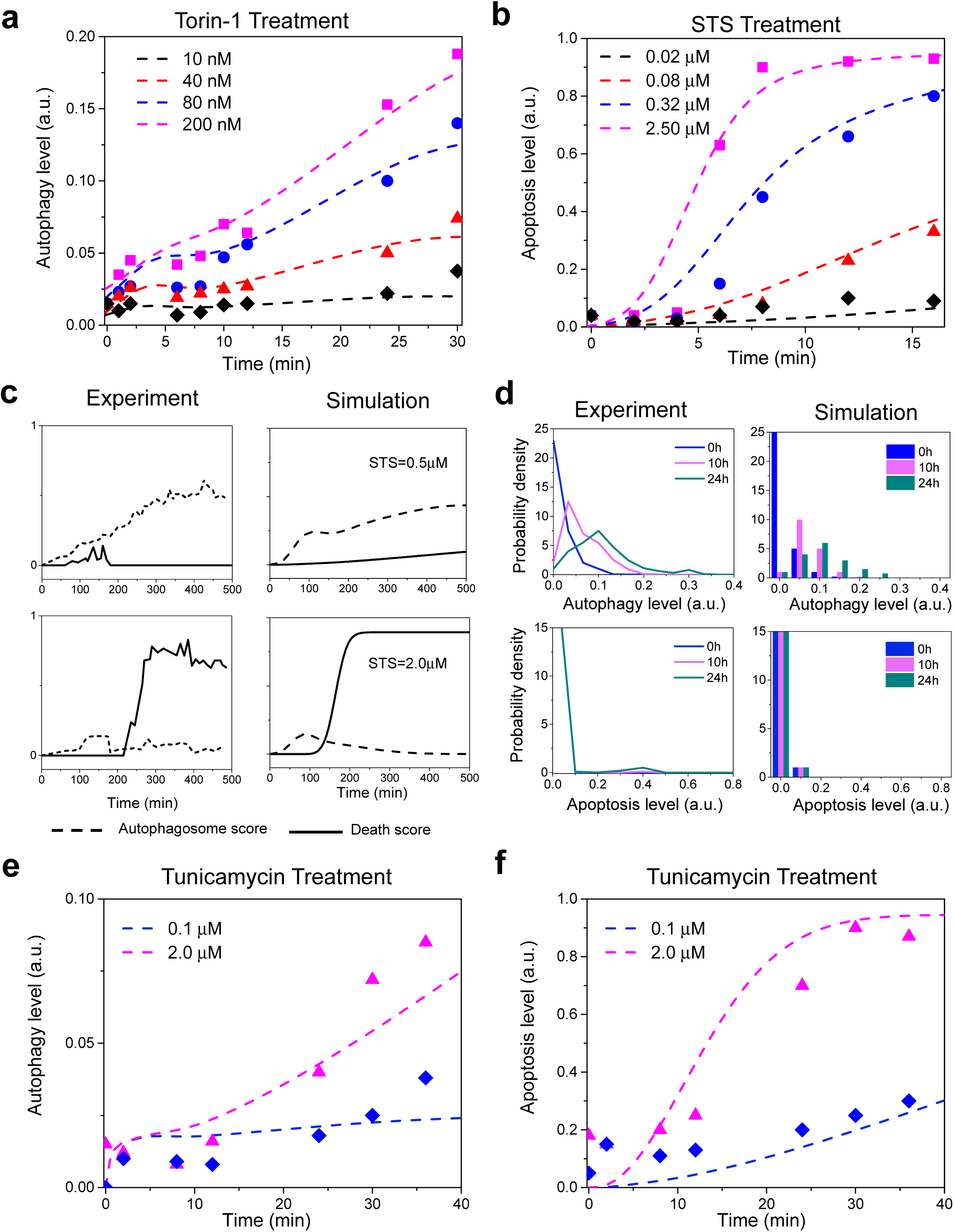
Model predictions and experimental validation. Experimental and simulated time evolution of autophagic and apoptotic response of individual cells to Torin-1 (**a**) and staurosporine (STS) treatment (**b**) and further comparison of autophagosome and death scores from experiments and simulations, in response to STS (**c**). Experimentally observed and computationally obtained kinetics of starvation-induced autophagy and apoptosis (**d**), and time courses of autophagy (**e**) and apoptosis (**f**) level in response to tunicamycin treatment. In panels **a**, **b**, **e** and **f**, the symbols designate the experimental data points.

We next proceed to the validation of our model. **Figure 2c** demonstrates that our model reproduces the differential dynamics of autophagy and apoptosis observed in the single-cell analysis^12^ (*the left panels* are adapted from Figure 5A-B of Xu et al, 2013). Specifically, STS-induced stress can stimulate autophagy (*dotted curve*) and/or apoptosis (*solid curve*) in the same individual cells, and autophagy precedes apoptosis. Under low stress (0.5 μM STS*; top diagrams*), the onset of autophagy (around 100 minutes) protects the cells from death. In contrast, under high-stress conditions (2.0 μM STS), temporary activation of autophagy is not sufficient to prevent apoptosis: the early autophagic response disappears with the cell commitment to apoptosis.

**Figure 2d** shows the comparison of the model-predicted histograms (*right panels*) with the experimentally observed probability densities (*left panels*) of autophagy and apoptosis levels in a population of H4 cells in response to 0, 10 and 24 h of starvation. 1,000 trajectories were generated in line with the prior distributions of initial concentrations. The simulations accurately reproduce the experimentally observed induction of autophagy but not apoptosis upon inhibition of mTOR, indicating that the model captures cell-to-cell variability.

We further validated our model using an additional test dataset^12^ on the time courses of autophagy and apoptosis of H4 cells in response to 0.1 and 2.0 μM tunicamycin (ER stress-inducer). **Figure 2e-f** shows that the model predictions (*dashed curves*) quantitatively reproduce the experimental data (*triangles and diamonds*) consistent with the observation that tunicamycin induces slower autophagy in H4 cells as compared with STS.

Model predictions quantitatively matched not only the training data (**Fig. 2a-b**) but also the test data (**Fig. 2c-f**). This increased our confidence that the model had not been overfitted and that it had adequate predictive power for the major effects regulating autophagy/apoptosis under different stress conditions.

### Sensitivity analysis indicates that Ca^2+^ release from ER, and regulation by p53, calpain, AMPK are key determinants of cell fate

We identified the components and reactions that are essential to cell fate decision using a multi-parametric sensitivity analysis (MPSA) based on SMC^11^ (see *Methods and Materials*). Model outputs used as criteria were the autophagy and apoptosis levels induced by 0.5 μM STS. **Figure 3a-b** show the global sensitivity values obtained by varying the kinetic parameters within the ranges listed in Supplementary **Table S3**.

**Figure 3.**
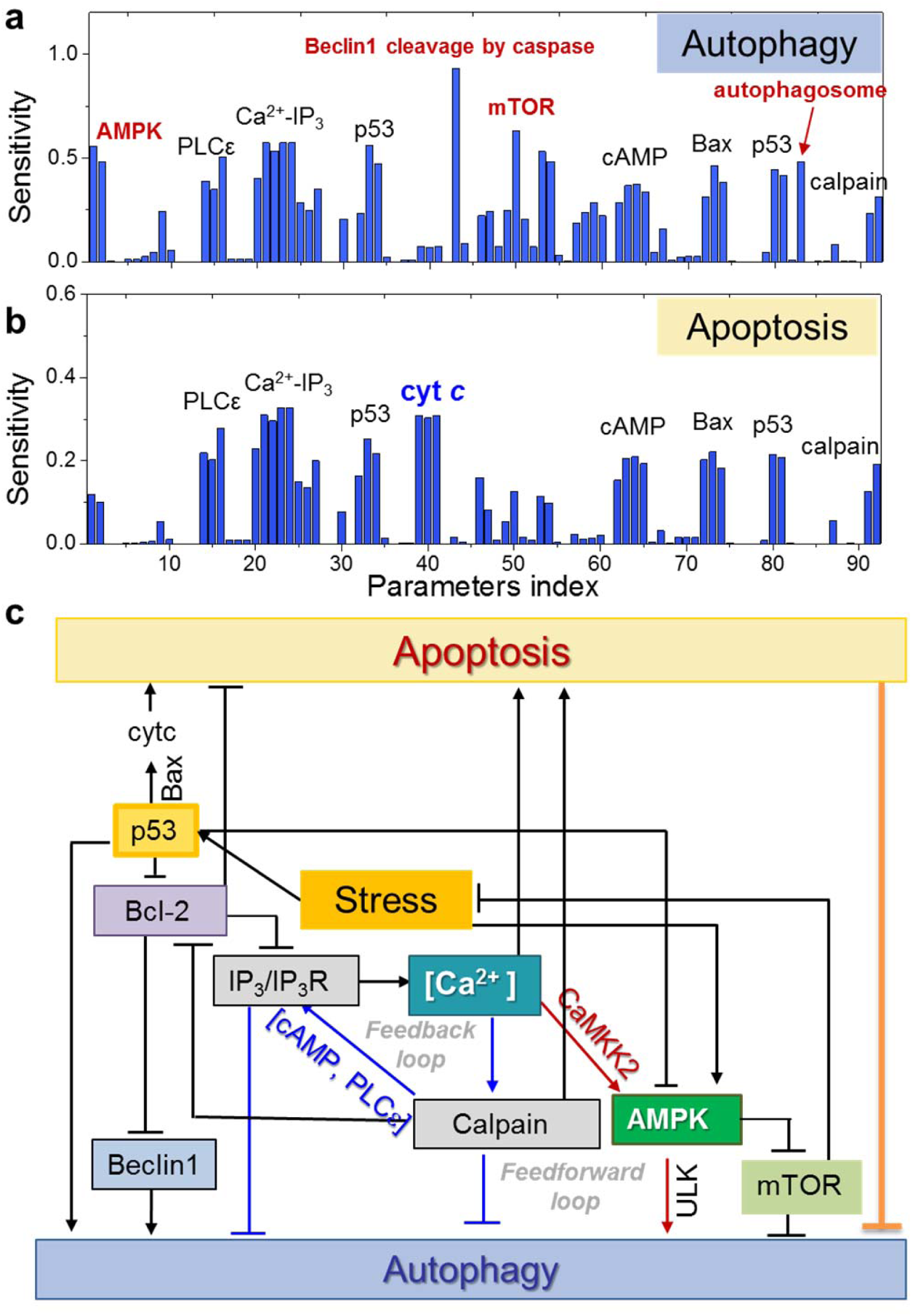
Sensitivity analysis. Peaks display the components distinguished by their strong effect on regulating autophagy (**a**) and apoptosis (**b**). Parameter index (abscissa) refers to Supplementary **Table S2**. (**c**) The core regulatory network composed of key determinants of cell decision.

Components whose kinetics exhibit strong effects are classified into three clusters: those sensitive to (i) both autophagy and apoptosis, (ii) only autophagy, and (iii) only apoptosis, indicated by labels colored *black*, *red* and *blue*, respectively. The reactions associated with cAMP-PLCε-IP_3_-Ca^2+^-calpain-pathway emerges as major determinants of both autophagy and apoptosis, pointing to the central role of [Ca^2+^(IC)]. p53 plays an important role in both apoptosis and autophagy, and couples to Ca^2+^ signaling via Bcl-2. Both Ca^2+^ and p53 regulate the activation/inhibition of AMPK, which, in turn, favors autophagy, hence the emergence of the AMPK peak in **Fig. 3a**. Notably, the cleavage of Beclin-1 by caspases is distinguished by its strong effect on autophagy (but not apoptosis). Beclin-1, like calpain, p53 and AMPK, occupies a central role in the crosstalk between the two pathways. The observed peaks thus underscore the significance of the crosstalk between these processes and also explain the biphasic behavior of autophagy observed in H4 cells in response to increasing STS dosage^12^. The cell’s first rescue response appears to induce autophagy at low stress/toxicity, and it resorts to apoptosis with increasing stress. The onset of apoptosis is accompanied by termination of autophagy, hence the observed modest control of autophagy by compounds known to dominate apoptosis.

**Figure 3c** summarizes the flow of information between key components distinguished in this section. The release of Ca^2+^ from the ER upon IP_3_-binding to IP_3_R is fueled by a positive feedback loop involving calpain* and PLCε*. The net effect of this loop is to suppress autophagy. However, this effect is countered by Bcl-2 that inhibits the IP_3_R. Yet, Bcl-2 simultaneously suppresses autophagy by inhibiting Beclin1; while calpain* and p53 moderate the effects of Bcl-2. The diagram also points to two competing roles of increased Ca^2+^: suppression of autophagy through calpain*; and promotion of autophagy, via a feedforward loop that involves CaMKKβ* and AMPK*. The autophagy-upregulating role of AMPK* is further reinforced by inhibition of mTORC1 and upregulaton of ULK1; however, AMPK* is deactivated by p53. These intricate network points to IP_3_/IP_3_R-regulated cytoplasmic Ca^2+^, p53, AMPK and calpain as master regulators of cell decision between apoptosis and autophagy.

### Cytoplasmic Ca^2+^ functions as a rheostat that fine-tunes the timing of autophagic and apoptotic responses

We next turn our attention to the mechanism of action of Ca^2+^(IC). The release of Ca^2+^ from the ER is enabled upon activation of ER membrane receptors by secondary messengers (represented here by IP_3_R activation by IP_3_). Ca^2+^ can conversely be transferred from the cytoplasm to the ER lumen by ATPase pumps such as sarco/ER Ca^2+^-ATPase (SERCA). To investigate the effect of alterations in [Ca^2+^(IC)] on autophagy regulation, we increased *in silico* the expression level of SERCA and simulated the autophagy profiles in response to low and high levels of stresses (Supplementary **Fig. S1**). Note that the same effect on [Ca^2+^(IC)] could alternatively be induced by inhibiting Ca^2+^ channels on cell membrane. Supplementary **Figure S1a** shows the decrease in [Ca^2+^(IC)] upon increasing [SERCA]_0_. At low stress, a graded increase in autophagic response is observed (**Fig. 4a**); whereas under high stress, the autophagic response is faster and of shorter duration: it reaches its peak around 10 h, even with moderate increases in [SERCA]_0_, after which it gives way to apoptosis (Supplementary **Fig. S1b**), i.e. the onset of apoptosis concurs with the weakening of autophagy. An abrupt surge in apoptotic response is robustly elicited by the initial reduction in [Ca^2+^(IC)] in accord with the bistability of apoptosis^31^, the transition being sharper and faster under high stress. Decrease in [Ca^2+^(IC)] under low stress, on the other hand, exerts a moderate effect on the strength and/or timing of apoptotic response (**Fig. 4b**).

**Figure 4.**
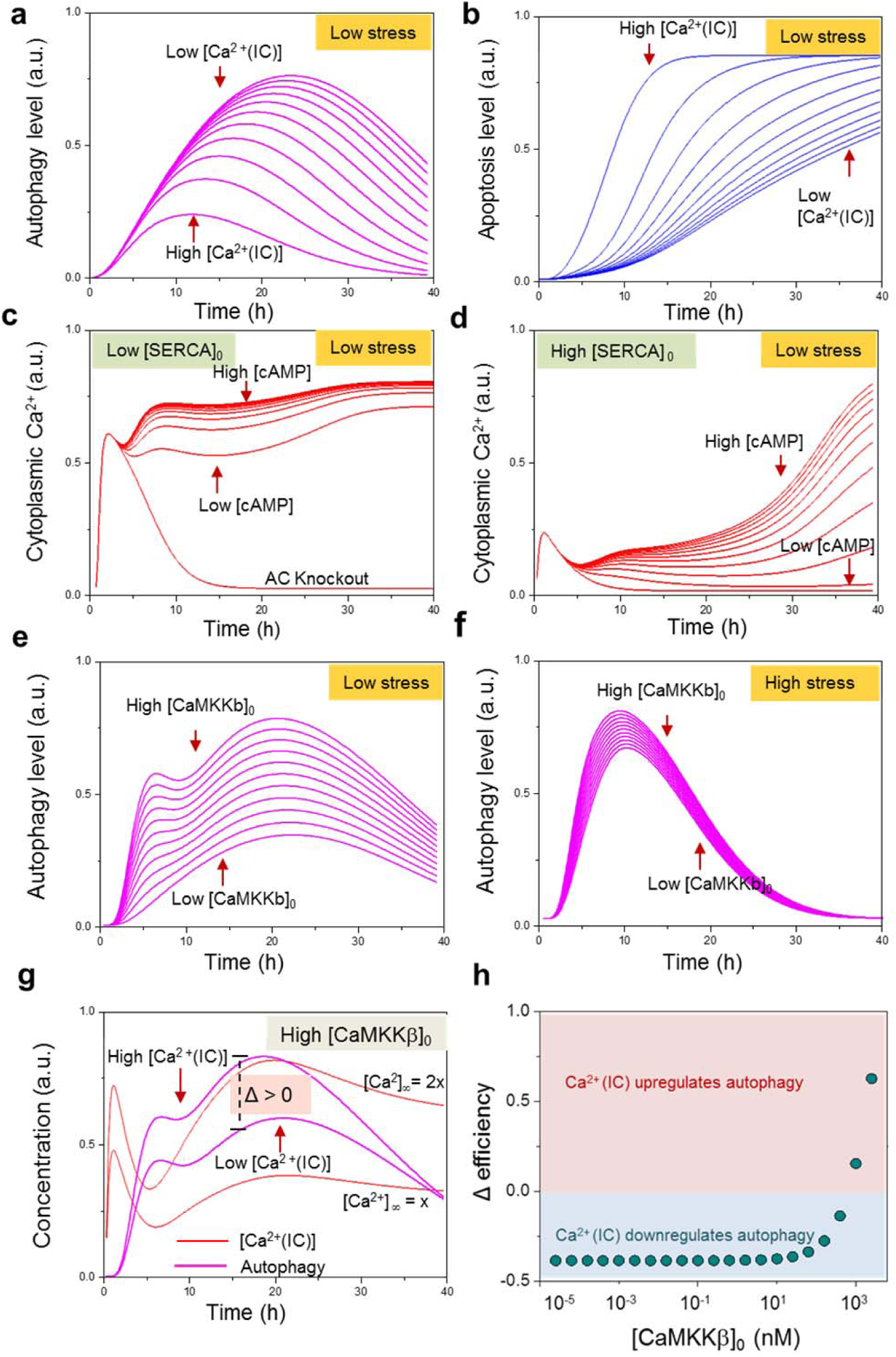
Role of cytoplasmic Ca^2+^ concentration in the onset of autophagy under low stress and its dependence on [CaMKKβ]. The stress is induced by administering a low dose (0.5 µM) of STS, except for panel **f** where [STS] = 2 µM. (**a**) Time evolution of autophagy as a function of cytoplasmic Ca^2+^ level (controlled by [SERCA]; **Fig. S1a**). (**b**) Apoptotic response under the same conditions. (**c-d**) Time evolution of [Ca^2+^(IC)] under different initial concentrations of AC, for low (**c**) and high (**d**) [SERCA]. [cAMP] produced by AC varies from 10 (low) to 100 nM (high). (**e-f**) Simulated development of autophagy for 0.01 < [CaMKKβ]_0_ < 1 nM, under low (**e**) and high (**f**) stress. The propensity of the cell for autophagy increases with increase in [CaMKKβ]_0_. (**g**) Simulated profiles of autophagy (*magenta*) accompanying the changes in [Ca^2+^(IC)] (*red*) in the presence of elevated [CaMKKβ]_0_. The enhancement in autophagy level due to change in [Ca^2+^(IC)]_∞_ by a factor of 2, is designated by ∆, which is maximum difference between the two curves. (**f**) Dose-response curve of ∆ as a function of [CaMKKβ]_0_. Intracellular Ca^2+^ downregulates autophagy in general (see **Fig. 4a** and Supplementary **S1b**) despite the opposing effect of CaMKKβ, except for elevated (> 10^3^nM) levels of CaMKKβ.

Taken together, our results suggest a ‘rheostat’ mechanism regulated by calcium signaling, which enables the cell to fine tune its response to stress. The cell can cope with low stress conditions by initiating an autophagic response, if [Ca^2+^(IC)] is sufficiently low, but if the stress is elevated, the same mediators of cell response that otherwise favor autophagy alter the cell commitment toward programmed death.

### Gα signaling and ensuing PLCε activation maintain cytoplasmic Ca^2+^ level through a positive feedback loop

As illustrated in **Fig. 1a** and summarized in **Fig. 3c**, [Ca^2+^(IC)] is regulated by a positive feedback loop formed G-protein α-subunit that drives cAMP production by AC, and IP_3_ signaling, which in turn activates calpain, which further stimulates the inositol pathway, and so on. Here we focus on this effect. To this aim, we varied [AC]_0_ and evaluated the effect of cAMP production and ensuing stimulation of EPAC and PLCε on [Ca^2+^(IC)]. Simulations were repeated under high and low influx of Ca^2+^ to the cytoplasm from other sources, here modulated by varying SERCA levels/activity. Supplementary **Figure S2a** shows that knocking down AC leads to a graded reduction of cAMP level as expected, the reduction being more pronounced with high [SERCA]_0_ (100nM). Of interest is the concurrent non-uniform changes in [Ca^2+^(IC)] (**Fig. 4c-d**). [Ca^2+^(IC)] exhibits a complex time evolution, depending on the extent of downregulation of EPAC/PLCε and upregulation of SERCA: (i) when the supply of Ca^2+^ is sufficiently high (e.g. low [SERCA]_0_ = 10nM), [Ca^2+^(IC)] is bistable; it maintains a high level even with a small stimulation of EPAC/PLCε activation via G-protein signaling, but it is severely depleted in the absence of such signaling (**Fig. 4c**), (ii) in the opposite case of an upregulated SERCA which promotes the influx of Ca^2+^(IC) into the ER, the suppression of [Ca^2+^(IC)] cannot be sustained due to the restoring effect Gα-signaling (**Fig. 4d**). The positive feedback loop mediated by calpain and PLCε activation thus plays a key role in restoring and maintaining the physiological cytoplasmic Ca^2+^ levels. Once Ca^2+^(IC) is boosted to a certain level, it robustly maintains its level by this cellular feedback.

### CaMKKβ shapes the role of cytoplasmic Ca^2+^ in regulating autophagy

Cytoplasmic Ca^2+^ plays a dual role in autophagy: it inhibits autophagy upon activation of calpain and IP_3_R; and enhances autophagy upon activation of AMPK in a CaMKKβ-dependent manner (respective *blue* and *red arrows* in **Fig. 3c**). This forms an incoherent type 1 feedforward loop (I1-FFL), a motif frequently seen in biological networks^32^. The effect of cytoplasmic Ca^2+^ thus depends on the balance between these opposing actions. Which effect dominates, under which conditions? **Figure 4a** suggest the dominance of the former (at least in H4 cells under low stress, with [SERCA]0 varying in the range 1-100nM), i.e. increase in [Ca^2+^(IC)] suppresses autophagy. To investigate whether there might be a reversal in this effect, we increased the amount of activated CaMKKβ. **Figure 4e** obtained under the same conditions with [SERCA]_0_ = 77nM, confirms that higher [CaMKKβ]_0_, which also leads to higher [CaMKKβ*], enhances autophagy. This effect is, however, temporary; it disappears after 20-40 hours, as autophagic response gives way to apoptosis. The termination of autophagy is expedited (with minimal dependence on [CaMKKβ]) under high stress (**Fig. 4f**).

To evaluate the overall role of cytoplasmic Ca^2+^ in regulating autophagy, we define a variable Δ that measures the *change in autophagy level* in response to doubling the steady state level [Ca^2+^(IC)]_∞_ of cytoplasmic Ca^2+^. As shown in **Fig. 4g**, for high [CaMKKβ]_0_, doubling [Ca^2+^(IC)]_∞_ (e.g. by knocking down SERCA) enhances autophagy (Δ > 0). In contrast, for low [CaMKKβ]_0_, doubling [Ca^2+^(IC)]_∞_ results in a decrease in autophagy (Δ < 0) (Supplementary **Fig. S3**).

This analysis implies that whether cytoplasmic Ca^2+^ up- or down-regulates autophagy depends on the initial concentration of CaMKKβ. **Figure 4h** shows the response curve of Δ as a function of [CaMKKβ]_0_, suggesting that cell types with different levels of CaMKKβ expression may exhibit opposite responses to autophagy-modulating drugs (e.g. verapamil) that target [Ca^2+^(IC)].

### *In silico* simulations reveal pharmacological strategies for controlling cell fate

The model developed here permits us to interrogate various treatment scenarios and evaluate their efficacies for either enhancing autophagy to protect normal cells or enhancing apoptosis to kill cancer cells. **Figure 5** illustrates the simulated treatment efficacy of low dose of eight FDA-approved drugs under three different levels of stress. The known (major) target and action of each drug are indicated. Odd and even columns represent the predicted fold changes in autophagy and apoptosis levels, respectively, in response to drug treatment.

**Figure 5.**
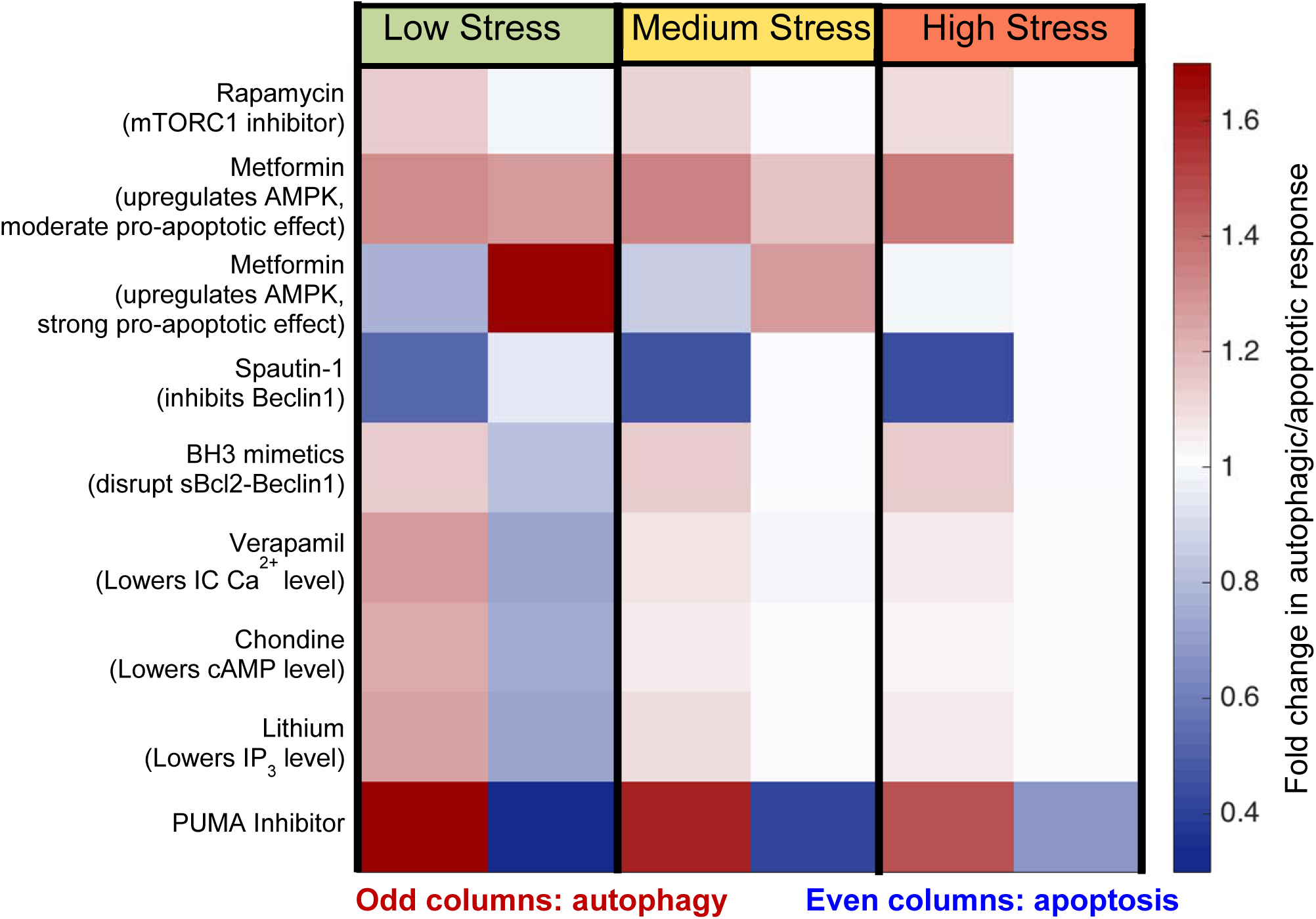
Simulated efficacy of pharmacological strategies. The simulated autophagy and apoptosis level in response to low, medium, and high dose of STS stress. The color-coded entries represent the fold change in autophatic (odd columns) and apoptotic (even columns) responses, relative to those in the absence of treatment.

The mTORC1 inhibitor rapamycin enhances autophagy irrespective of the stress level, and has practically no effect on apoptosis. This is consistent with experimental observations using H4 cell line (neuroglioma)^33^.

AMPK was distinguished here as a key up-regulator of autophagy, and upregulating AMPK enhances autophagy and may slightly hinders apoptosis due the elimination of the stress by autophagy (Supplementary **Fig. S4a**). Metformin is known to upregulate AMPK, while a recent study also indicate that metformin downregulates the expression of Bcl-2 and upregulates the expression of Bax^34^. Inclusion of these promiscuous effects of metformin in the model, led to either simultaneous enhancement of autophagy and apoptosis or enhanced apoptosis but suppressed autophagy, depending on the strength of pro-apoptotic effects of metformin on a particular cell line (**Fig. 6**). These results are consistent with the observations that metformin promote autophagy and apoptosis in esophageal squamous cell carcinoma^35^, while promoting apoptosis but suppressing autophagy in glucose-deprived H4IIE hepatocellular carcinoma cells^36^. The model thus permits us to evaluate the effects of promiscuous drugs, or polypharmacological effects.

Spautin-1, an inhibitor of Beclin-1, effectively suppresses autophagy under all conditions and its effect on apoptosis is minimal, except for low stress conditions. This is consistent with the experimental observations in H4 cells and Madin-Darby canine kidney epithelia (MDCK) cells^37^.

BH3 mimetics (which disrupts the interaction between Bcl-2 and Beclin-1), verapamil (which lowers [Ca^2+^(IC)]), clonidine (which lowers [cAMP]), and lithium (which lowers [IP_3_]) have similar actions and efficacies. Under low level of stress, they enhance autophagy and suppress apoptosis, while for medium and high levels of stress, they essentially enhance autophagy. BH3 mimetics (e.g.ABT737) have been reported to induce autophagy for multiple cell lines (e.g. HeLa, U2OS, HCT116)^38,39^, suggesting that the interaction between Bcl-2 and Beclin-1 might be a robust drug target for enhancing autophagy. Both verapamil and clonidine have been shown to induce autophagy in PC12 cells^40^. Our prediction is also consistent with the observation that with the same dosage (1µM), verapamil induces a higher level of autophagy than does clonidine. The role of lithium on inducing autophagy has been demonstrated in H4 cells and PC12 cells^41,42^.

Interestingly, PUMA inhibitor shows remarkable pro-survival effects: it enhances autophagy and suppresses apoptosis under all conditions. This is consistent with the results from sensitivity analysis which highlighted the strong inhibitory effect of apoptosis on autophagy (via the cleavage of Beclin-1 by caspases). It implies that radiation mitigators may enhance pro-survival autophagy in injured cells to further prevent cell death. PUMA inhibitors have been shown to efficiently inhibit apoptosis using multiple cell lines^43^, while their upregulation of autophagy requires further confirmation.

Taken together, these results indicate that: (i) to protect normal cells against stress-induced cell death, upregulation of AMPK, which simultaneously activates multiple downstream autophagic pathways, and inhibition of PUMA^43^, could be highly efficacious strategies; (ii) In contrast, Beclin-1 inhibitors such as spautin-1 may improve the efficacy of apoptosis-inducers and may be advantageously used for pre-empting autophagic response and enabling the elimination of cancer cells with high apoptotic thresholds.

## Discussion

Autophagy has been viewed as a form of programmed cell death in several studies, named as ‘autophagic cell death’ (ACD) or ‘type II cell-death’, since programmed cell death was often associated with enhanced autophagy and depended on autophagy-associated proteins to a certain context^44^. However, recent evidence led to a debate on the existence of ACD^9,45^. Our current understanding is that though ACD exists in rare situations, a more general scenario is that autophagy precedes apoptosis in the same cell in order to adapt to or cope with non-lethal stress^1^ and apoptosis occurs when the stress exceeds a critical threshold of intensity or duration^8^. The present study provides a firm quantitative description of the validity of this interplay between autophagy and apoptosis. In most cases, autophagy and apoptosis mutually inhibit each other and the onsets of autophagy and apoptosis are governed by cell fate decision processes that have broad pathophysiological implications^8,46^. We presented here a calibrated computational model for the kinetics of (macro)autophagy and apoptosis pathway crosstalk along with several testable results.

Our analysis identified a core regulatory network for autophagic and apoptotic responses (**Fig. 3c**). Specifically, cytoplasmic Ca^2+^, p53, AMPK, calpain and Bcl-2 emerged as key components whose expression/activities significantly affect cell decision. Ca^2+^ and p53 act as master regulators that tightly control cell decisions through AMPK and Bax activation pathways, respectively. A positive feedback loop formed by Gα signaling and inositol pathways is critical to maintaining [Ca^2+^(IC)], which in turn governs the graded autophagic response through a feedforward loop mediated by CaMKKβ and calpain. Our model also predicts that the CaMKKβ activation may act as a determinant of the dual/opposite roles of cytoplasmic Ca^2+^ as well as treatment efficacy. While increased [Ca^2+^(IC)] generally inhibits autophagy (in favor of apoptosis), this effect can be reversed in cells that express high levels of CaMKKβ.

Previous modeling works have focused on apoptosis^47-49^. Recent progress on modeling of autophagy either focused on specific modules such as autophagic vesicle dynamics^50^ and mTORC1-ULK1 interaction^51^ or used over-simplified networks^52,53^. In a recent study^54^, a relatively larger cancer-specific model of 13 ODEs has been built and calibrated. Unfortunately, it has not been further advanced and thus no new knowledge has been obtained so far.

Here, we have developed a first comprehensive and calibrated kinetic model of the autophagy-apoptosis crosstalk. Our model consists of 94 species which cover many important stress-sensing pathways (**Fig. 1b**). The model is generic and its parameters can be calibrated to capture the pathway activation/dynamics in specific cell types. We have demonstrated that our model was able not only to reproduce single-cell and population-based measurements of autophagic and apoptotic responses of neuroglioma cells under nutritional, genotoxic, and ER stresses (**Fig. 2**), but also to serve as a platform for gaining insights into the underlying time-dependent interactions (**Fig. 3-4**) as well as interrogating potential therapeutic strategies (**Fig. 5**).

Live imaging and single-cell analysis have indicated that autophagy proceeds via a graded dynamics whereas apoptosis onset obeys a switch-like behavior^12^. However, the underlying mechanism remained unclear. Our previous work^13^ have shown that the positive feedback loop, Bax*→caspase→tBid→Bax*, is critical for sustained caspase activity and thereby the switch-like dynamics of apoptosis. In the context of the autophagy-apoptosis crosstalk, this positive feedback loop is also responsible for the all-or-none bimodal dynamics of apoptosis, since it mediates the pro-apoptotic roles of other species including calpain and Ca^2+^(IC). As autophagy precedes apoptosis, it could start eliminating dysfunctional entities when the causing stress levels are not sufficiently high to trigger the caspase/tBid feedback loop. Autophagic events thus delay, if not prevent, apoptosis (maintaining ‘off’ state). This points to the important timing of intervention for pre-empting the potential commitment of the cell to apoptosis, while the cell is dealing with dysfunctional elements via autophagy. Our model allows for interrogating the network of interactions and cell fate decision in the presence of one or more pharmacological interventions and assessing effective treatments that favor autophagy (for healthy cells under stress) or apoptosis (for cancer cells) under different stress conditions, as illustrated in **Fig. 5**.

When the stress level or duration reaches an apoptotic threshold, the positive feedback loop that ensures the sustained caspase cascade and the apoptotic machinery switches from ‘off’ to ‘on’ state. Upon committing to apoptosis, the cell shuts down autophagy and recruits many players (otherwise involved in both autophagic and apoptotic events) to promote apoptosis. Previous work^52^ hypothesized that this change in course is mainly due to the cleavage of Beclin-1 by caspase. Our sensitivity analysis corroborates that Beclin-1 cleavage by caspases is an event of utmost importance (**Fig. 3a**). However, our sensitivity analysis also highlights other important players/interactions. In particular, a unique incoherent type 1 feedforward loop (I1-FFL) involving CaMKKβ, AMPK, calpain, cAMP and IP_3_R emerged here as a major determinant of [Ca^2+^(IC)], and consequently up- or down-regulation of autophagy. A dual role of Ca^2+^(IC) in autophagy emerges from a large number of contradicting evidences (see review^28^). Elevated levels of Ca^2+^ have been reported to promote autophagy, while inhibitors of intracellular Ca^2+^ currents have been found to promote autophagy^55^. Our results indicated that the balance between the two (*red* and *blue*) branches in **Fig. 3c** defines the decision/fate of the cell. Calpain is part of the cAMP-mediated positive feedback loop for sustaining [Ca^2+^(IC)]; it also activates apoptosis via Bid activation. Thus, its complex role hinders calpain as a modulator to shape the role of Ca^2+^(IC). Controlling CaMKKβ levels on the other hand emerges as a viable therapeutic strategy.

CaMKKβ is a versatile regulator of the CaMKs and involved in regulating many cellular processes such as glucose homeostasis and inflammation^29^. The expression of CaMKKβ varies in different cell types and tissues. Our results implicated that overexpression of CaMKKβ can switch cytoplasmic Ca^2+^ from an inhibitor to an enhancer of autophagy (**Fig. 4h**). Ca^2+^(IC)-modulating drugs (e.g. verapamil) should thus be used in a cell/context-dependent manner, since the expression and activity level of CaMKKβ might influence the drug efficacy. Recent evidence showed CaMKKβ is expressed at very low levels in normal prostate, but accumulates in prostate cancer cells^56^. Consequently, enhanced activation of the CaMKKβ-AMPK pathway may increase autophagy and elevate the threshold for the onset of apoptosis, and thereby assist in the survival of prostate cancer cells. In such context, Ca^2+^(IC)-reducing drugs might be potentially used in combination with apoptosis-inducers to suppress autophagy and further enhance apoptosis.

Calcium is released from the ER by ryanodine receptors too^57^. The opening of ryanodine receptors is triggered by ADP ribose. This mechanism is similar to the IP_3_R-dependent calcium release, thus in our model, IP_3_R may be viewed as a membrane protein representative of ligand-gated receptors that release calcium from the ER, including RYR. Further, sources of Ca^2+^ intake from the EC region include voltage-gated calcium channels and ligand-gated receptors such as NMDA receptors; and PMCA (and Na^+^/Ca^2+^ exchangers) pump Ca^2+^ from the cytosol to the EC region^58^. In the current model, we implicitly modeled these processes by assuming constant concentration level of Ca^2+^ intake from the EC medium, but these proteins actually represent alternative targets for modulating autophagic responses via altering Ca^2+^ levels. Extension of the current model to explicitly include these components and their interactions will increase the utility of the model as a platform for designing and evaluating alternative treatment strategies.

### Methods

The details of our methods for mathematical modeling, parameter estimation, and sensitivity analysis are presented in Supplementary **SI Methods**. The reactions in **Fig. 1B** and associated parameters are presented in Supplementary **Table S2**.

## Acknowledgement

This work was funded by the NIH grants P01DK096990, U19AI068021, and P41GM103712.

## Author contributions

BL and IB conceived and designed research; BL performed research; BL and IB analyzed the results with input from ZNO; BL, ZNO, HB, GAS, GSP, DHP and IB wrote the paper. All authors approved the final version of the paper.

